# Olfactory cortex output pathways exhibit robust differences in their ability to drive learned aversion

**DOI:** 10.1101/2023.06.04.541965

**Authors:** Inês Vieira, Dmitri Bryzgalov

## Abstract

The olfactory (piriform) cortex contains a large diversity of neural cell types, defined by their molecular identity, morphology, functional properties and projection target specificity. How these distinct neural cell types contribute to the control of olfactory-driven behaviors remains unknown. Here, we use a behavioral task in which optogenetic activation of subpopulations of piriform neurons is associated with an unconditioned aversive stimulus. We find that piriform neurons projecting to the olfactory bulb or the medial prefrontal cortex efficiently drive a learned escape behavior. In contrast, neurons projecting to the cortical amygdala or the lateral entorhinal cortex failed to support a behavioral response. Our results suggest that different piriform neural subnetworks exhibit robust differences in their ability to relate neural activity to behavioral output.

## Introduction

The detection of odorants is accomplished by odorant receptors, expressed on the dendrites of olfactory sensory neurons in the main olfactory epithelium. In mice, olfactory sensory neuron express one of 1,000 odorant receptor genes present in the genome (Zhang and Firestein, 2002; Niimura, 2012), and project to one of two spatially stereotyped glomeruli in the olfactory bulb (OB) (Mombaerts et al., 1996; Ressler et al., 1994; Vassar et al., 1994). The output neurons of the olfactory bulb, the mitral and tufted cells, then send projections through the lateral olfactory tract to several higher olfactory centers in the brain, including the piriform cortex, the anterior olfactory nucleus, the olfactory tubercle (OT), the cortical amygdala (CoA), and the lateral entorhinal cortex (lENT) (Haberly and Price, 1978; Ghosh et al., 2011; Sosulski et al., 2011).

The piriform cortex is the largest cortical area receiving direct inputs from OB mitral and tufted cells, mitral and tufted cells projections to piriform are widespread and diffuse, lacking apparent topographical organization (Ghosh et al., 2011; Igarashi et al., 2012; Sosulski et al., 2011). Single piriform neurons receive convergent inputs from mitral and tufted cells belonging to multiple glomeruli, allowing for the integration of the segregated patterns of odor-evoked glomerular activity (Davison and Ehlers, 2011; Franks et al., 2011). Furthermore, piriform networks are shaped by olfactory learning and experience, and optogenetic activation of piriform neural ensembles is sufficient to drive learned behaviors (Chapuis and Wilson, 2011; Choi et al., 2011). Together, these data suggest that the piriform cortex plays important functions in odor perception, and olfactory learning and memory.

Odor information encoded by piriform ensembles must then be transmitted to downstream target areas to support multimodal sensory integration and motor control. Piriform cortex neurons send feedback projections to the OB as well as efferent projections to several cortical and sub-cortical targets, including the medial prefrontal cortex (mPFC), the OT, the CoA, and the lENT (Chen et al., 2014; Diodato et al., 2016; Haberly and Price, 1978; Johnson et al., 2000; Mazo et al., 2017). Recent experiments have begun to shed light onto the organization of piriform output pathways. For example, piriform neurons projecting to different subdivisions of the orbitofrontal cortex exhibit distinct spatial topography along the antero-posterior axis of piriform cortex (Chen et al., 2014). Furthermore, it has recently been shown that distinct subpopulations of piriform projection neurons segregate into distinct piriform layers: neurons projecting to the OB and the mPFC are enriched in deep piriform layers IIb and III, while neurons projecting to the CoA and the lENT are predominantly present in the superficial piriform layer IIa. These different classes of piriform projection neurons can also be distinguished based on their morphology and the expression of distinct molecular markers (Diodato et al., 2016). However, the relevance of this organization of distinct piriform output pathways for olfactory-driven behaviors remains unknown.

The functional characterization of distinct piriform output pathways, through manipulation of neural activity, is complicated by the fact that OB mitral and tufted cells project to multiple higher olfactory centers in cortex, which, in turn, are strongly interconnected (Franks et al., 2011; Illig and Haberly, 2003; Johnson et al., 2000; Luskin and Price, 1983; Mazo et al., 2017). Such parallel neural pathways may interfere with perturbations in the activity of subpopulations of piriform neurons, potentially confounding the interpretations of behavioral phenotypes. We therefore devised an alternative experimental strategy to probe the functional properties of distinct piriform subpopulations, which is based on the direct optogenetic activation of piriform neurons and allows us to test the sufficiency of distinct piriform subpopulations to drive a behavioral response. We find that subpopulations of projection neurons in the deep piriform layers IIb and III, selectively targeted based on their connections with the OB or the mPFC, reliably drive a learned escape behavior, upon association of light-induced neural activity with an unconditioned aversive stimulus. In contrast, neurons in superficial piriform layer IIa, targeted based on their connections with the CoA or the lENT failed to support this learned escape behavior under our experimental conditions. Our data provide evidence for differences in the ability of piriform subnetworks to link neural activity with behavioral output.

## Results

### OB- and lENT-projecting piriform neurons have distinct output targets in the anterior telencephalon

To confirm and extend the characterization of the projection targets of distinct classes of piriform neurons we used an intersectional viral tracing approach. We injected the retrogradely transported Canine Adeno Virus 2 expressing Cre recombinase (CAV2-Cre) (Junyent and Kremer, 2015; Schwarz et al., 2015) into the OB and the lENT of adult mice, and we infected piriform cortex neurons with conditional Adeno-Associated Virus expressing channelrhodopsin (AAV-ChR2-EYFP). Cre-mediated recombination in piriform neurons results in the expression of ChR2-EYFP in distinct subpopulations of piriform projection neurons. ChR2-EYFP is robustly expressed on cell soma and axon terminals, and thus provides an anterograde neural tracer to identify brain regions innervated by piriform fibers. As previously reported, we observed that OB- and lENT-projecting piriform neurons can be distinguished based on their laminar positioning and morphology (Diodato et al., 2016 **Figure 1b, d** and data not shown). We then analyzed EYFP fluorescence on axonal processes in coronal sections through the brain. As expected, in mice in which CAV2-Cre was injected into the OB, we detected dense labeling of EYFP-positive axons in the granule cell layer of the OB (**Figure 1e**). In addition, we observed EYFP-positive axons in the mPFC, the OT, and the insular cortex (IC) (**Figure 1f**). These data suggest that OB-projecting neurons send axon collaterals to several other target areas in the anterior telencephalon. In contrast, targeting lENT-projecting neurons resulted in only minimal labeling of the OB, the mPFC, the OT, and the IC (**Figure 1h, i**). Labeling could be observed in superficial fiber tracts in the anterior telencephalon, however, the precise projection targets of these fibers could not be resolved in this analysis. We also observed differences in the more posterior projection targets of piriform neurons targeted by CAV2-Cre injections into the OB and the lENT (**Figure 1g, j**). However, the potential spread of virus into adjacent areas including the CoA and the lENT precluded a more detailed analysis. Taken together, these experiments demonstrate that distinct piriform projection neurons can be targeted using intersectional viral-genetic tools, and that OB- and lENT-projecting piriform neurons have largely non-overlapping target areas.

**Figure 1.**
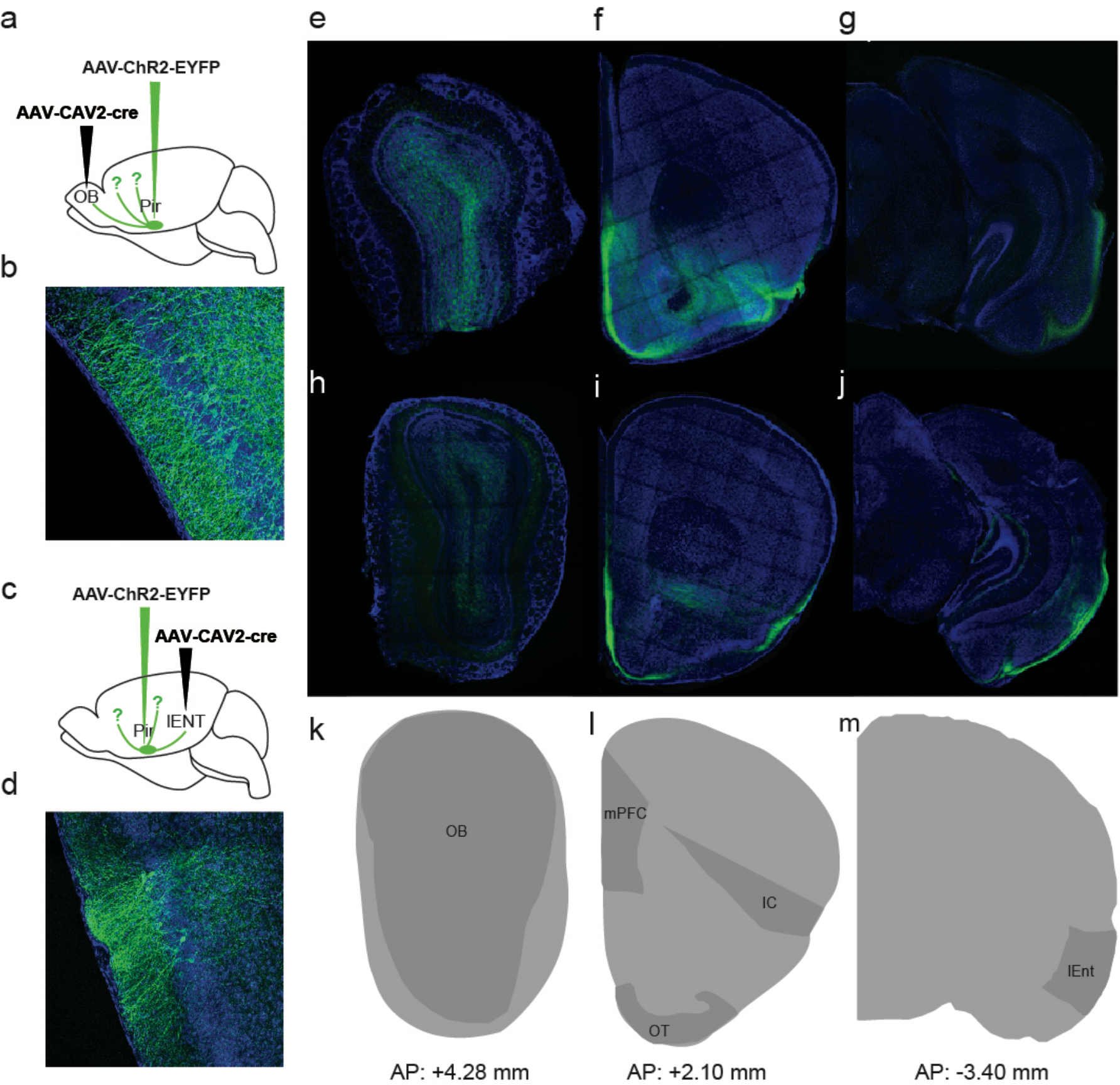
Target areas of OB- and lENT-projecting piriform neurons. Schematic representations of the injection sites of conditional AAV-ChR2-EYFP in the piriform, and CAV2-Cre virus in the OB (a) and in the lEnt (c), below is represented the piriform injection site (b,d). Coronal sections of the target areas of OB -projecting piriform neurons (e-g) and of I Ent -projecting piriform neurons (h-j) to the olfactory bulb (OB), medial prefrontal cortex (mPFC),insular cortex (IC), olfactory tubercle (OT) and the lateral entorhinal cortex (lEnt). EYFP labeling in green, neurotrace counterstain in blue, (k-m) In dark grey, are represented the main target regions of piriform neurons; below the reference AP coordinates of each coronal section, n (number of mice) = 4, 4 sections per mouse.

### Photostimulation of OB- and CoA-projecting piriform neurons activates distinct subnetworks of neurons

We next photoactivated distinct ChR2-expressing subpopulations of piriform projection neurons and monitored light-induced neural activity using immunohistochemical detection of the transcription factor c-Fos. Stimulation through implanted optical fibers in piriform cortex resulted in robust induction of c-Fos immunoreactivity (**Figure 2a, c**). While the numbers of c-Fos-positive neurons were highly variable (number of Fos neurons following photostimulation of OB-projecting neurons, average per mouse: 6356, 4433, 114, and 161 and number of Fos neurons following photostimulation of CoA projecting neurons, average per mouse: 665, 926, 1003, 480), likely due to the variability of the numbers of infected neurons and the efficiency of their activation by light, the laminar distribution of c-Fos-positive neurons was consistently different between the two targeted subpopulations. When photostimulating OB-projecting neurons, c-Fos-positive cells were predominantly observed in the deep piriform layers IIb and III (**Figure 2b**). In contrast, when stimulating CoA-projecting neurons, c-Fos expression was robustly enriched in superficial piriform layer IIa cells (**Figure 2d**).

**Figure 2.**
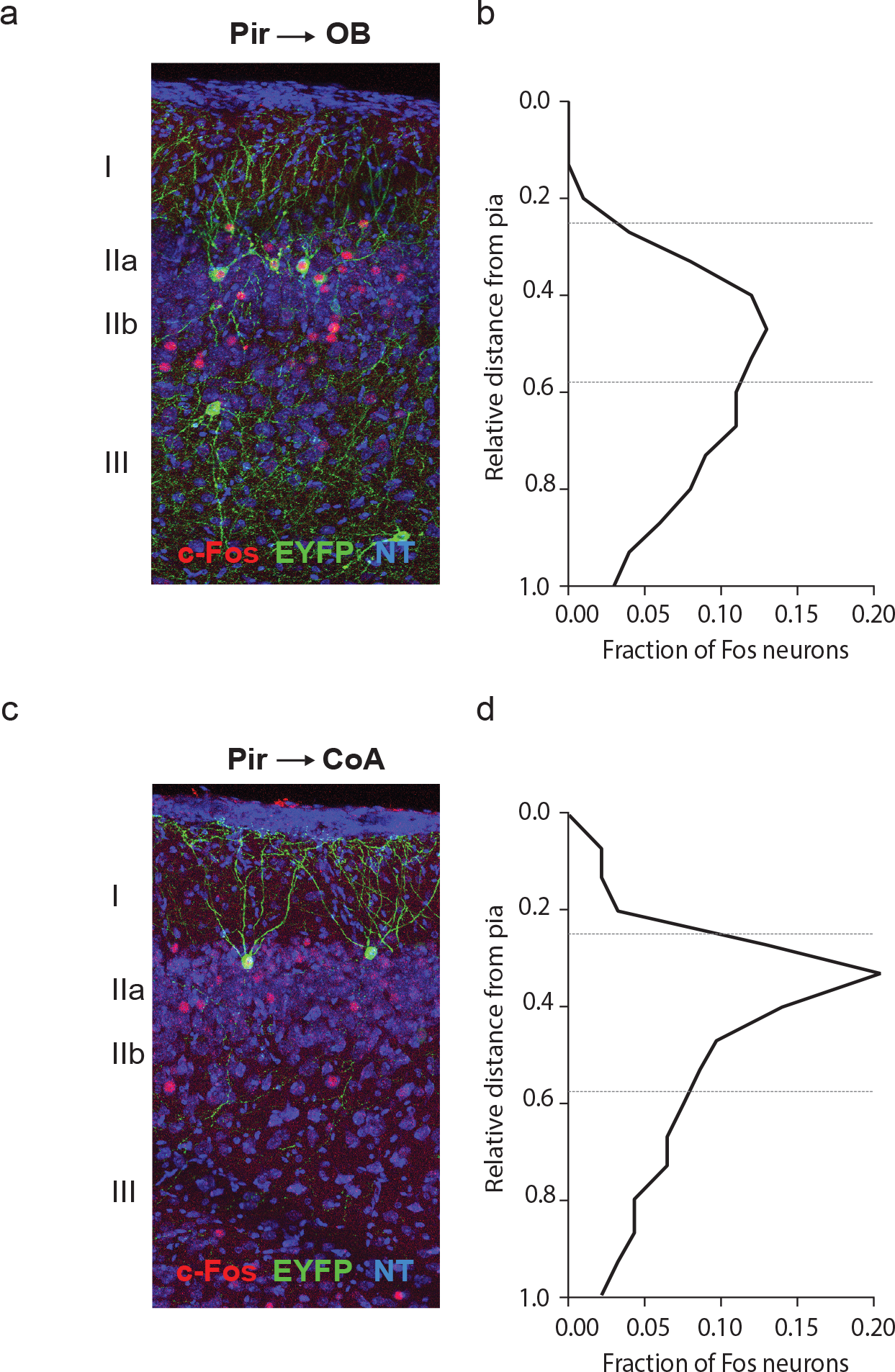
Photoactivation of distinct subpopulations of piriform projection neurons. Immunohistochemical analysis of coronal sections of piriform cortex after photostimulation of subpo-pulations of ChR2-expressing neurons. AAV-ChR2-EYFP-expressing neurons are in green, c-Fos immunoreactivity in red, neurotrace (NT) in blue. (**a**) Subpopulations of neurons targeted via their projections to the olfactory bulb (OB). (**c**) Subpopulations of neurons targeted via their projections to the cortical amygdala (CoA). (**b**,**d**) Laminar distribution of c-Fos-expressing neurons upon photosti-mulation of OB- and CoA-targeted subpopulations. c-Fos-expressing neurons are enriched in the superficial layer IIa in CoA-targeted subpopulations, while c-Fos expression is observed in deeper piriform neurons in OB-targeted mice. n (number of mice) = 4.

It is important to note that light-induced c-Fos expression was not confined to ChR2-expressing piriform neurons. c-Fos-positive, ChR2-negative neurons were observed within and outside of the targeted sublayer of piriform cortex, i.e. layers IIb and III for OB-projecting neurons, and layer IIa for CoA-projecting neurons. The spread of c-Fos immunoreactivity beyond the neurons directly responsive to photostimulation is consistent with the activation of piriform neurons through extensive recurrent excitatory connections (Franks et al., 2011; Poo and Isaacson, 2011). Nevertheless, despite such intra-cortical propagation of light-induced neural activity, photoactivation of OB- and CoA-projecting neurons resulted in the activation of distinct ensembles of piriform subnetworks.

### Different classes of piriform projection neurons differ in their ability to drive learned aversion

We next adapted a previously established behavioral paradigm, in which photoactivation of piriform neurons paired with foot shock drives a learned escape behavior (Choi et al., 2011). To test whether the activity of distinct subpopulations of piriform neurons was sufficient to elicit conditioned aversion, we selectively expressed ChR2 in OB- and CoA-projecting neurons. An additional cohort of mice, in which we expressed the calcium indicator GCaMP3 instead of ChR2 in piriform neurons, was used to control for the specificity of light-induced neural activity as the conditioned stimulus (CS). Training was performed in a rectangular box in which mice could move freely. Foot shock as the unconditioned stimulus (US) was applied to the side of the box where the mouse was located at the time of photostimulation, allowing the mouse to escape from the aversive stimulus by running towards the opposite side of the box. The CS-US presentation was randomly applied to either side, depending on the location of the mouse. After two training sessions, mice were tested in a box with identical dimensions, to determine whether photoactivation of neurons alone was sufficient to elicit learned aversion.

Escape behavior was quantified using three parameters. We measured the maximum speed of mice and the distance traveled, during a 10 second (s) time window before and after the beginning of photostimulation. Furthermore, we determined the time between the beginning of photostimulation and a behavioral response (“reaction time”) during the same time period. For each mouse, data were averaged across 7 individual trials. Similar results were obtained when analyzing individual trials independently (Supplementary Figure 1). We found that photostimulation of OB-projecting neurons reliably elicited a robust behavioral response. Mice exhibited an increase in the maximum speed after photostimulation, an increase in the distance travelled, and a short reaction time (maximum speed before vs. after photostimulation: 19.5 vs. 30.6 cm/s, p = 0,0098; distance run before before vs. after photostimulation: 35.7 vs. 44.4 cm, p = 0,2324; reaction time: 4.2 s; **Figure 3c-e**). In contrast, photoactivation of CoA-projecting neurons failed to produce a behavioral response (maximum speed before vs. after photostimulation: 18.3 vs. 17.8 cm/s, p =0,5887; distance run before vs. after photostimulation: 31.2 vs. 28.9 cm, p = 0,4961 ; reaction time: 6,7 s; **Figure 3c-e**). A similar lack of behavioral response was evident in control mice, in which GCaMP3 instead of ChR2 was expressed in piriform neurons (maximum speed before vs. after photostimulation: 10.53 vs. 8.7 cm/s, p = 0,4403; distance run before vs. after photostimulation: 16.3 vs. 15.6 cm, p > 0,99, reaction time: 8.5 s; **Figure 3c-e**).

**Figure 3.**
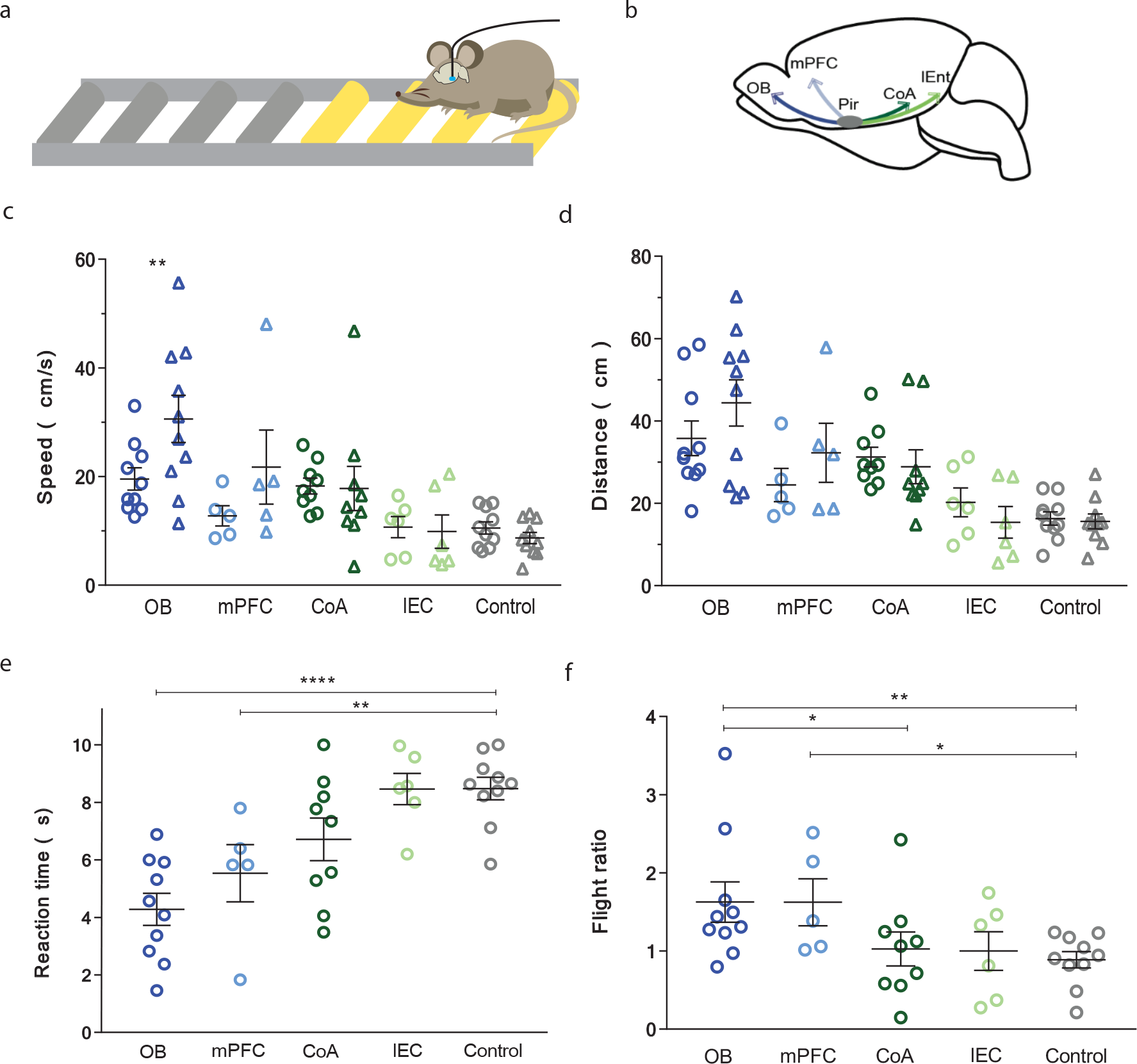
OB- and mPFC-projecting, but not CoA- and lENT-projecting piriform neurons are sufficient to drive learned aversion. (**a**) Fear conditioning assay set up and optogenetics in freely moving mice. (**b**) Schematic representation of the injection sites of AAV-flex-ChR2 in the Piriform and CAV2-Cre in the different target areas: OB, mPFC, CoA and lEnt. (**c**) Maximum speed in cm/s, measured during 10 s windows before and after photostimulation, of mice in which distinct subpopulations of piriform neurons were entrained to drive a learned escape behavior. OB-projecting neurons: n = 10, CoA-projecting neurons: n = 9, mPFC-projecting neurons: n = 5, lENT-projecting neu-rons: n = 6, and AAV-GCamp3 expressing neurons (control group), n = 10. (**d**) Distance traveled before and after photostimulation. (**e**) Reaction time after photostimulation. (**f**) Flight ratio, defined as the maximum speed after photostimulation, divided by the maximum speed before photostimulation. Each dot represents data obtained from individual mice, averaged across seven trials. Bars represent the mean +/-SEM.

To further extend our analysis we next targeted mPFC- and lENT-projecting piriform neurons. mPFC-projecting neurons substantially overlap with OB-projecting neurons, while lENT-projecting neurons largely overlap with CoA-projecting neurons (Diodato et al., 2016). Based on the observations described above, we predicted that mPFC-projecting neurons, similar to OB-projecting neurons, may support a conditioned behavioral response, while lENT-projecting neurons, similar to CoA-projecting neurons, may fail to do so. We found that photoactivation of mPFC-projecting neurons was indeed sufficient to elicit learned aversion, reflected in an increase in the maximum speed and distance traveled, and a short reaction time (maximum speed before vs. after photostimulation: 12.76 vs. 21.75 cm/s, p = 0,0625; distance run before vs. after photostimulation: 24.5 vs. 32.3 cm, p = 0,313, reaction time: 5.5 s, **Figure 3c-e**). In contrast, photoactivation of lENT-projecting neurons failed to produce a reliable behavioral response (maximum speed before vs. after photostimulation: 10.68 vs. 9.87 cm/s, p = 0,5887; distance run before vs. after photostimulation: 20.2 vs. 15.4 cm, p = 0,3095, reaction time: 8.5 s, **Figure 3c-e**). Finally, to compare between different cohorts of mice, which can exhibit differences in baseline motor activity, we calculated the escape (flight) ratio, defined by the maximum speed after photostimulation divided by the maximum speed before photostimulation (**Figure 3f**). This analysis further illustrates that OB- and mPFC-projecting subpopulations of piriform neurons are sufficient to drive a learned escape behavior, while CoA- and lENT-projecting neurons fail to do so. Taken together, these data identify functional differences in the ability of distinct piriform subnetworks to relate neural activity to behavioral output.

### Behavioral output does not correlate with the number of active neurons

Differences in the ability of distinct subpopulations of piriform neurons to drive learned aversion may reflect differences in the numbers of photoactivated neurons, rather than functional differences between different piriform subnetworks. To test this possibility, we trained an additional cohort of mice, and we determined the numbers of c-Fos-positive neurons directly after behavioral testing. Consistent with our results described above, we observed higher escape ratios in mice in which OB-and mPFC-projecting neurons had been targeted, compared to two mice in which CoA-projecting neurons had been targeted (one mouse with OB-targeted neurons: escape ratio 1.49, one mouse with mPFC-targeted neurons: escape ratio 2.14, two mice with CoA-targeted neurons: escape ratios 1.25 and 1.11). However, we found fewer c-Fos-expressing neurons in the OB- and mPFC-targeted mouse compared to the two CoA-targeted mice (mouse with OB-targeted neurons: 373 c-Fos- positive piriform cells, mouse with mPFC-targeted neurons: 241 c-Fos-positive cells; mice with CoA-targeted neurons: 599 and 412 c-Fos-positive cells). Thus, a higher number of photoactivated, c-Fos-positive neurons does not correlate with an increase in learned aversion, suggesting that functional differences in the different neural subnetworks underlie the observed differences in the behavioral responses.

## Discussion

We established a viral-genetic approach to target distinct subpopulations of piriform projection neurons, and we tested their ability to relate neural activity to behavioral output. We found that neurons in the deep piriform layers IIb and III were capable of eliciting robust escape behavior upon pairing of light-induced neural activity with foot shock as the unconditioned stimulus. In contrast, targeting superficial piriform layer IIa cells failed to produce a behavioral response under these experimental conditions. Our data thus identify functional differences amongst different piriform output pathways.

We previously showed that OB- and mPFC-projecting piriform neurons are enriched in piriform layers IIb and III, exhibit the morphological characteristics of superficial and deep pyramidal cells and express molecular markers that delineate their projection target specificity (Diodato et al., 2016). In contrast, the majority of CoA- and lENT-projecting neurons are piriform semilunar cells and are located in the superficial piriform layer IIa. Previous work further suggests that piriform projection neurons can extend axon collaterals to multiple target regions (Diodato et al., 2016; Mazo et al., 2017). Neural tracing experiments using the retrogradely transported CAV2 virus provide an opportunity to label the entire axonal projections of neurons, based on a single selected output target area (Schwarz et al., 2015). We found that OB-projecting neurons elaborate axons to multiple additional target areas, including the mPFC, IC, and the OT. In contrast, axons emanating from lENT-projecting neurons only minimally innervated these brain regions. An important limitation of this experimental approach is the likely spread of conditional AAV-ChR2-EYFP virus into areas adjacent to piriform cortex. While viral infection was well confined within piriform layers and across its dorso-ventral axis, viral spread appeared to be less confined along the antero-posterior axis. We observed a few infected neural cell bodies in regions posterior to piriform cortex, such as in the amygdala and entorhinal cortex (data not shown). Therefore, our analysis of axonal projection patterns is potentially confounded by labeled axons from nearby regions, in particular in more posterior brain regions. Future experiments using sparse and focal expression of a neural tracer, combined with the tracing of individual axons from identified neurons in cleared brain preparations should provide improved resolution to resolve these questions.

Photostimulation of ChR2-expressing piriform neurons resulted in c-Fos immunoreactivity in ChR2-positive and ChR2-negative neurons, both within and beyond the targeted piriform sub-layers (Figure 2). Such propagation of light-induced neural activity is likely to reflect the activity of excitatory connections within and across piriform layers (Franks et al., 2011; Suzuki and Bekkers, 2011). However, despite extensive intracortical connectivity, photostimulation of distinct ChR2-expressing subpopulation of piriform neurons resulted in the activation of piriform ensembles with distinct laminar distribution. It will be interesting to explore patterns of light-induced neural activity beyond the piriform cortex. The analysis of c-Fos expression patterns in direct piriform target areas as well as in more distant neural processing centers may provide insights into how neural activity is propagated through neural circuits in the brain to elicit motor output.

Photoactivation of OB- and mPFC-projecting piriform neurons effectively drove learned aversion. Two training sessions, each consisting of 10 pairings of photoactivation with foot shock, were sufficient to elicit a robust escape behavior when photoactivtion was presented in the absence of foot shock during testing. In contrast, photoactivation of CoA- and lENT-projecting neurons did not produce a behavioral response under the same experimental conditions. A trivial explanation for this observation could be that viral injections into the CoA and the lENT did not target a sufficiently large population of piriform neurons. We consider this explanation unlikely, based on the following two observations. First, the number of light-induced c-Fos-positive neurons was highly variable when targeting OB-projecting neurons (Figure 2). Despite such variability, we found that OB-targeted neurons reliably elicited a robust escape behavior. Second, we tested, albeit in a small cohort of mice, whether the number of c-Fos-positive neurons correlated with escape behavior. We found that fewer than 400 c-Fos-positive neurons in OB- and mPFC-targeted mice were sufficient to elicit conditioned aversion, while more than 400 c-Fos-positive neurons in CoA-targeted mice failed to elicit the behavioral response. A more plausible explanation of the observed differences is that distinct piriform subpopulations require a different photostimulation regimen to entrain a learned behavior. For example, photostimulation of CoA- or lENT-projecting neurons at different frequencies and/or laser power may support the association of foot shock with a behavioral response. Such a model would suggest that functional differences between the piriform subnetworks, such as differences in neural excitability and network plasticity underlie the observed differences in behavioral output.

An alternative possibility consistent with our observations is that distinct subnetworks of piriform neurons are specialized to control different behavioral responses. Fear conditioning in mice, for example, can result in escape behavior or freezing, depending on experimental constraints (Fadok et al., 2017). It will therefore be interesting to test if in conditions in which mice cannot escape from the unconditioned stimulus, CoA- and lENT-projecting neurons can support a conditioned freezing response. Finally, different subpopulations of piriform neurons may be dedicated to support behaviors of different valence, such as aversive or appetitive conditioning, and experience-dependent social behaviors (Choi et al., 2011). The experimental approach we have developed provides an opportunity to probe the sufficiency of different piriform output pathways to support different learned behaviors.

## Methods

### Mice

Adult (8- to 12-week-old) C57BL6/J male wild-type mice were used in this study. Mice were housed at the animal facility at the CIRB, Collège de France. All experiments were performed according to European and French National institutional animal care guidelines (protocol number B750512/00615.02).

### Immunohistochemistry

Mice were deeply anaesthetized with pentobarbital and transcardially perfused with 20 ml of PBS, followed by 10 ml of 4% paraformaldehyde. Brains were post-fixed for 4 h in 4% paraformaldehyde at 4 °C. Coronal sections (200 μm thick) were prepared using a vibrating-blade microtome (Microm Microtech). Sections were rinsed in PBS and permeabilized in PBS/0.1% Triton X-100 for 1 h, and blocked in PBS/0.1% Triton X- 100/2% heat-inactivated horse serum (Sigma) for 1 h. After incubation with primary antibodies at 4 °C overnight, sections were rinsed in PBS/0.1% Triton X-100, three times for 20 min at room temperature, blocked in PBS/0.1% Triton X-100/2% heat-inactivated horse serum for 1 h and incubated with secondary antibodies overnight at 4 °C. The following antibodies were used at the indicated dilutions: rabbit anti-cfos 1:500 (Santa Cruz sc-7270), and chicken anti-GFP 1:1000 (Abcam ab13970). Appropriate secondary antibodies (1:1000) conjugated to Cy3 (Jackson Labs) or Alexa 488 (Molecular Probes) were incubated together with Neurotrace counterstain (1:500, Invitrogen). Sections were mounted on SuperFrost Plus (Menzel-Gläser) microscope slides in Fluorescent Vectashield Mounting Medium (Vector). Images were acquired with a Leica SP5 confocal microscope, with 10x or 20x objectives.

### c-Fos stimulation protocol

Animals injected with CAV2 virus expressing Cre-recombinase in the region of interest (CAV2-Cre), and conditional AAV vectors carrying channelrhodopsin (ChR2-eYFP) were photostimulated (20Hz, 30sec per minute for 10minutes) and processed for immunohistochemical analysis one hour later.

### Quantification of c-Fos positive neurons

Immunohistochemistry was performed on piriform sections obtained 0.8–1.2 mm posterior to bregma. The percentage of neurons expressing c-Fos at the center of the injection site (3 histological sections) was obtained by manual counting. All image processing and quantification was performed in Fiji and Adobe Photoshop CS5. A given field of view was divided into 15 bins, and the fraction of cells in each bin was calculated as the total number of c-Fos+ cells in each bin divided by the total number of cells in the field of view.

### Stereotaxic viral injections and fiber implantation

Mice were anaesthetised intraperitoneally with ketamine/xylazine (100 and 20 mg per kg of body weight, respectively) and prepared for surgery in a stereotactic frame (David Kopf Instruments). For viral injections, a small craniotomy was made above the injection site. For anterograde neural tracing experiments and behavior experiments, 0.7 μl of AAV5.hSyn.hChR2 (H134R)-eYFP.WPRE.hGHpA was stereotaxically injected into the piriform cortex. In the control group, 0.6 μl of AAV1.hSyn.GCamp3-eYFP.WPRE.hGHpA was injected in the piriform cortex. 0.3 μl of canine adenovirus (CAV)-2-Cre was injected into the OB, the CoA, the mPFC or the lEnt. Virus was injected using a glass pipette with a 10–20 μm tip diameter. AAVs were obtained from the University of Pennsylvania (Penn Vectors). CAV2-Cre was obtained from the Montpellier Vector Platform (PVM). The following coordinates, based on the Paxinos and Franklin Mouse Brain Atlas were used: piriform cortex: anterior-posterior (AP) -0.60 mm, medio-lateral (ML) 3.95 mm, dorso-ventral (DV) -3.97 mm; OB: AP 0.75 mm and ML -0.75 mm coordinates from the midline rhinal fissure, DV -0.70 mm; medial PFC (including the prelimbic (PrL), infralimbic (IL), and cingulate (Cg) cortex): AP 0.54mm, ML 0.36mm, DV -1.70mm; posteromedial cortical amygdaloid nucleus: AP -2.80, ML 2.76, DV -4.8; lENT: AP -3.90, ML 3.8, DV 2.90.

For behavior experiments, optical fiber implantations were performed directly after viral injection; the skull was cleaned and covered with a layer of Super Bond C and B (Phymep). An optical fiber (200µm, 0.22 NA) housed inside a connectorized implant (SMR, Doric Lenses) was inserted into the brain, with the fiber tip positioned 200µm above the infection target (piriform cortex). The implants were secured with dental acrylic (Pi-Ku-Plast HP 36, Bredent). Before being moved back to their cages, after the surgery, mice were put in a warm chamber at 37C until full recovery.

### Aversive behavior paradigm

Photostimulation protocol All animals were single-housed and kept on a 12h/12h day/night cycle. Behavioral tests began 13 days post-surgery, and were carried out 6 hr after onset of the day period. Behavioral conditioning took place over 5 days. On the first day, mice were habituated to the fear conditioning box for 40 min, on the following day and on the fourth day, the mice were trained to associate the photostimulation with the foot shock and on the last day mice were tested, 24 hr after the second training session.

The optical fiber implant was connected to a mono fiber patch cord (MFP_200/240/900-0.22_FC-SMC, Doric Lenses) that was connected to a fiberoptic rotary joint (FRJ_1×2i_FC-2FC_0.22, Doric Lenses), which receives the light that passes through galvanometric mirrors from the free*-*space DPSS laser beam (MBL-III-473, CNI lasers). Laser output was maintained at 7-10 mW as measured at the end of the fiber. The conditioning apparatus was a rectangular chamber (9 cm W x 57 cm L x 16 cm H) with a stainless-steel rod floor. Each half of the conditioning apparatus was connected to an electrically operated switch, which was connected to an aversive stimulator (115 V, 60 Hz, Med Associates) and to a microcontroller board (Mega 2560, Arduino), allowing foot-shock to be applied independently to either side. The testing apparatus was a rectangular chamber (9 cm W x 57 cm L x 16 cm H) with a white PVC floor. Before each training session, laser beam output intensity and shape were adjusted and electrical current in the rod floor was measured with an aversive stimulation current test package (Med Associates). Photostimulation and foot shock were controlled using Arduino 1.8.3 open software.

Experimental animals were allowed to habituate to the apparatus for 5 min. The conditioning paradigm consisted of 3s of photostimulation (20 Hz square-wave shape, 25ms pulses) followed immediately by a 0.5s, 0.65 mA foot shock. Foot shock was applied only when the animal was in or near either end of the apparatus, forcing the animal to run to the opposite side. Photo stimulation/shock pairings were spaced 3-4 min apart. Each of the two training sessions consisted of 10 photo simulation/foot shock pairings, for a total of 20 pairings. Photo stimulation was applied 7 times over the testing session, every 3-4 min. All sessions were video-recorded.

Videos recorded during the testing session were analyzed during the 10 seconds time period before and after stimuli presentation using a custom-written Matlab script to quantify: maximum speed before and after stimuli presentation, distance run before and after stimuli presentation and reaction time after stimuli presentation.

### Statistics

Statistical analysis was performed in R and GraphPad Prism 7 software. Behavioral responses per mouse and per trial for speed and distance were compared before and after CS+ presentation and analysed for statistical significance using the non-parametric Wilcoxon test. Behavioral responses in between mice and trials of different behavior conditions for reaction time and flight ratio were analysed for statistical significance using the non-parametric Mann-Whitney test. In Figure 3 and Supplementary Figure 1, p values ≤ 0.05 are represented with *, p value ≤ 0.01 are represented with **, p value ≤ 0.001 are represented with *** and p values ≤ 0.0001 are represented with ****.

## Acknowledgments

We thank A.F. for the help with conceptualization, original draft writing, supervision, and funding acquisition. We thank G. Dugué, Y. Dupraz and G. Paresys for their help with optogenetics and the setup of the behavior asset. We thank members of the Fleischmann lab for their input during scientific discussions. This work was supported by grants from the Fondation pour la Recherche Médicale AJE201106. I.V was supported by a PhD international fellowship (Ref. SFRH/BD/91460/2012) from the Foundation for Science and Technology, Portugal and D.B. was supported by a graduate scholarship from École des Neurosciences de Paris Île-de-France.

## Author contributions

Conceptualization, I.V. and A.F.; Experiments, Methodology and Analysis, I.V. and D.B.; Writing and Reviewing, A.F. and I.V.

