## Supplemental information for "Olfactory cortex output pathways exhibit robust differences in their ability to drive learned aversion"

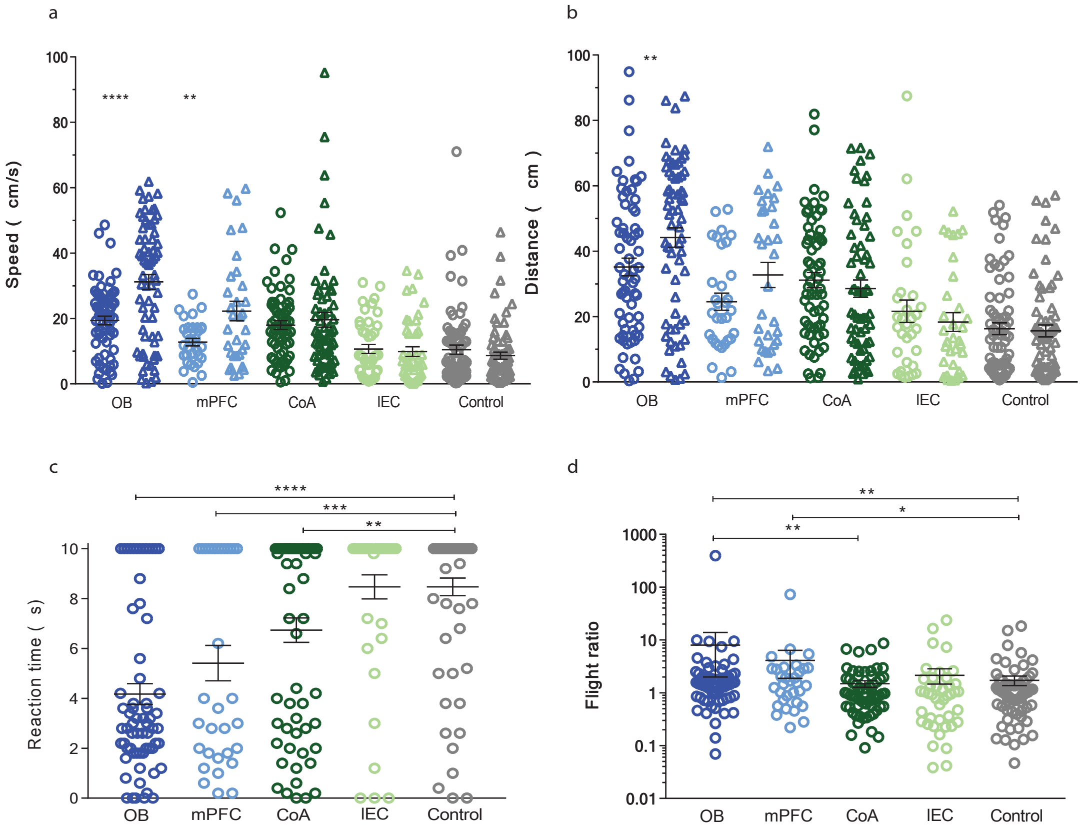


**Supplementary Figure 1.** OB- and mPFC-projecting, but not CoA- and lENT-projecting piriform neurons are sufficient to drive learned aversion, trial-by-trial data analysis. a) Maximum speed per trial in cm/s, measured during 10 s windows before and after photostimulation, of mice in which distinct subpopulations of piriform neurons were entrained to drive a learned escape behavior. OB-projecting neurons: n = 80, CoA-projecting neurons: n = 75, mPFC-projecting neurons: n = 32, lENT-projecting neurons: n = 42, and AAV-GCamp3 expressing neurons (control group), n = 69. (b) Distance traveled per trial before and after photostimulation. (c) Reaction time per trial after photostimulation. (d) Flight ratio, defined as the maximum speed after photostimulation, divided by the maximum speed before photostimulation per trial. Each dot represents individual trials. Bars represent the mean +/- SEM.
